# Avoiding false discoveries: Revisiting an Alzheimer’s disease snRNA-Seq dataset

**DOI:** 10.1101/2023.04.01.535040

**Authors:** Alan E Murphy, Nurun Nahar Fancy, Nathan G Skene

**Author notes:** Correspondence should be addressed to A.M. (email: Alan Murphy) and N.S. (email: Nathan Skene).

## Abstract

Mathys *et al*., conducted the first single-nucleus RNA-Seq study (snRNA-Seq) of Alzheimer’s disease (AD)^1^. The authors profiled the transcriptomes of approximately 80,000 cells from the prefrontal cortex, collected from 48 individuals – 24 of which presented with varying degrees of AD pathology. With bulk RNA-Seq, changes in gene expression across cell types can be lost, potentially masking the differentially expressed genes (DEGs) across different cell types. Through the use of single-cell techniques, the authors benefitted from increased resolution with the potential to uncover cell type-specific DEGs in AD for the first time^2^. However, there were limitations in both their data processing and quality control and their differential expression analysis. Here, we correct these issues and use best-practice approaches to snRNA-Seq differential expression, resulting 549 times fewer differentially expressed genes at a false discovery rate (FDR) of 0.05.

## Main

Mathys *et al*.’s^1^ data processing and quality control strategy for their single-nucleus RNA-Seq (snRNA-Seq) data was state-of-the-art at this time. Furthermore, the authors took extra measures in an attempt to ensure the reliability of their results. Our re-analysis is not a criticism of this but endeavours to raise awareness and provide recommendations for rigorous analysis of single-cell and single-nucleus RNA-Seq data (sc/snRNA-Seq) for future studies. Most importantly, we aim to ensure that the Alzheimer’s disease research field does not focus on spuriously identified genes. To this end, our questions of Mathys *et al*.,^1^ focus around their data processing and their differential expression analysis. Firstly, in relation to their processing approach, the authors discussed the high percentages of mitochondrial reads and low number of reads per cell present in their data. This is indicative of low cell quality^3^ however, we believe the authors’ quality control (QC) approach may not capture all of these low quality cells. Moreover, the authors did not integrate the cells from different individuals to account for batch effects. As the field has matured since the author’s work was published, dataset integration has become a common step in single-cell RNA-Seq protocols and is recommended to remove confounding sources of variation^4,5,6^. To gain advantage of these recent approaches, we used scFlow^7^ to reprocess the author’s data. This pipeline included the removal of empty droplets, nuclei with low read counts and doublets followed by embedding and integration of cells from separate samples and cell typing. ScFlow combines best practice approaches for processing sc/snRNA-Seq datasets, see Supplementary Note 1 for a detailed explanation of these steps. Re-processing resulted in 50,831 cells passing QC, approximately 20,000 less the authors’ post-processing set with differing cell type proportions (Supplementary Fig. 1).

In regards to data quality, it is worth noting that over 99% of nuclei had less than 200 genes expressed (Supplementary Table 3). While this QC step was not unique to our reprocessing, the authors made the same exclusion in their analysis^1^, it highlights the relatively low quality of the data which may be attributable to the early stage of snRNA-Seq technology of the time. For example, Brase *et al*.’s recent study of snRNA-Seq of autosomal dominant AD^8^ used a more stringent cut-off for the minimum number of expressed genes and still kept 27% (122 times more) of the assayed cells after all QC steps. Moreover, as mentioned, the authors discussed the high percentages of mitochondrial reads in their data. The differences in approaches to filtering based on the proportion mitochondrial reads accounts for the notable discrepancy in the number of nuclei after QC between our approach and the authors. Our approach used a 10% cut-off for the proportion of mitochondrial reads in a nuclei, as set out in Amezquita *et al*.’s best-practice guidelines^5^, which is less stringent than Seurat’s guidelines (5%)^9^ or that from Heumos *et al*. (8% from a median absolute deviations based cut-off selection)^4^. Conversely, the authors filtered out high mitochondrial read nuclei based on clusters from their t-SNE projection of the data^1^. Even at our lenient cut-off, over 16,000 nuclei that were removed in our QC pipeline were kept by the author’s (Supplementary Fig. 2), explaining the discrepancy in the number of nuclei after QC. Based on Supplementary Fig. 2, it is clear the author’s approach was ineffective at removing nuclei with high proportions of mitochondrial reads which is indicative of cell death^3,4^. We have made the data from our alternative processing approach publicly available (through Synapse: https://doi.org/10.7303/syn51758062.1) so researchers can utilise this resource free of low quality nuclei.

Our second question of Mathys *et al*.,^1^ is their differential expression approach. The authors conducted a differential expression analysis between the controls and the patients with AD pathology, concentrating on six neuronal and glial cell types; excitatory neurons, inhibitory neurons, astrocytes, microglia, oligodendrocytes and oligodendrocyte precursor cells, derived from the Allen Brain Atlas^10^. They performed downstream analysis on their identified DEGs and investigated some of the most compelling genes in more detail. Therefore, all findings put forward by their paper were based upon the validity of their differential expression approach. However, for this approach, the authors conducted a two-part, cell and patient level analysis. The cell-level analysis took each cell as an independent replicate and the results of which were compared for consistency in directionality and rank of their DEGs against their patient level analysis, a Poisson mixed model. The authors identified 1,031 DEGs using this combinatorial approach – DEGs requiring an FDR <0.01 in the cell-level and an FDR<0.05 in the patient level analysis. It is important to note that this cell-level differential expression approach, also known as pseudoreplication, over-estimates the confidence in DEGs due to the statistical dependence between cells from the same patient not being considered^11,12,13,14^. When we inspect all DEGs identified at an FDR of 0.05 from the authors’ cell-level analysis, this number increases to 14,274. Pseudobulk differential expression (DE) analysis has recently been proven to give optimal performance compared to both mixed models and pseudoreplication approaches^11,12,15,16^. It aggregates counts to individuals thus accounting for the dependence between an individual’s cells.

Here, to compare the effect of the different DE approaches in isolation, we apply a pseudobulk DE approach, sum aggregation and edgeR LRT^17^, to the authors original processed data. We found 26 unique DEGs when considering the six cell types used by the authors (Supplementary Table 2, Supplementary Fig. 3). This was 549 times fewer DEGs than that reported originally at an FDR of 0.05. When we compare these DEGs, we can see that the absolute log2 fold change (LFC) of our DEGs is 15 times larger than the authors’; median LFC of 2.34 and 0.16, despite the authors’ DEGs having an FDR score 8,000 times smaller; median FDR of 2.89x10^-7^ and 0.002 (Fig. 1a-b). Although we examined a high correlation in the genes’ fold change values across our pseudobulk analysis and the author’s pseudoreplication analysis (Pearson R of 0.87 for an adjusted p-value of 0.05, Supplementary Table 3), the p-values and resulting DEGs vary considerably. The correspondence in fold change values is expected given the approaches are applied to the same dataset whereas the probabilities, which pertain to the likelihood that a gene’s expressional changes is related to the case/control differences in AD, importantly do not align. We can show that this stark contrast is just an artefact of the authors taking cells as independent replicates and thus artificially inflating confidence by considering the Pearson correlation between the number of DEGs found and the cell counts (Fig. 1c-e). There is a near perfect, positive correlation between DEG and cell counts for the authors’ pseudoreplication analysis (Fig. 1c) and for the 1,031 genes from the authors’ combinatorial approach (Fig. 1d) which is not present in our pseudobulk re-analysis (Fig. 1e).

**Fig 1:**
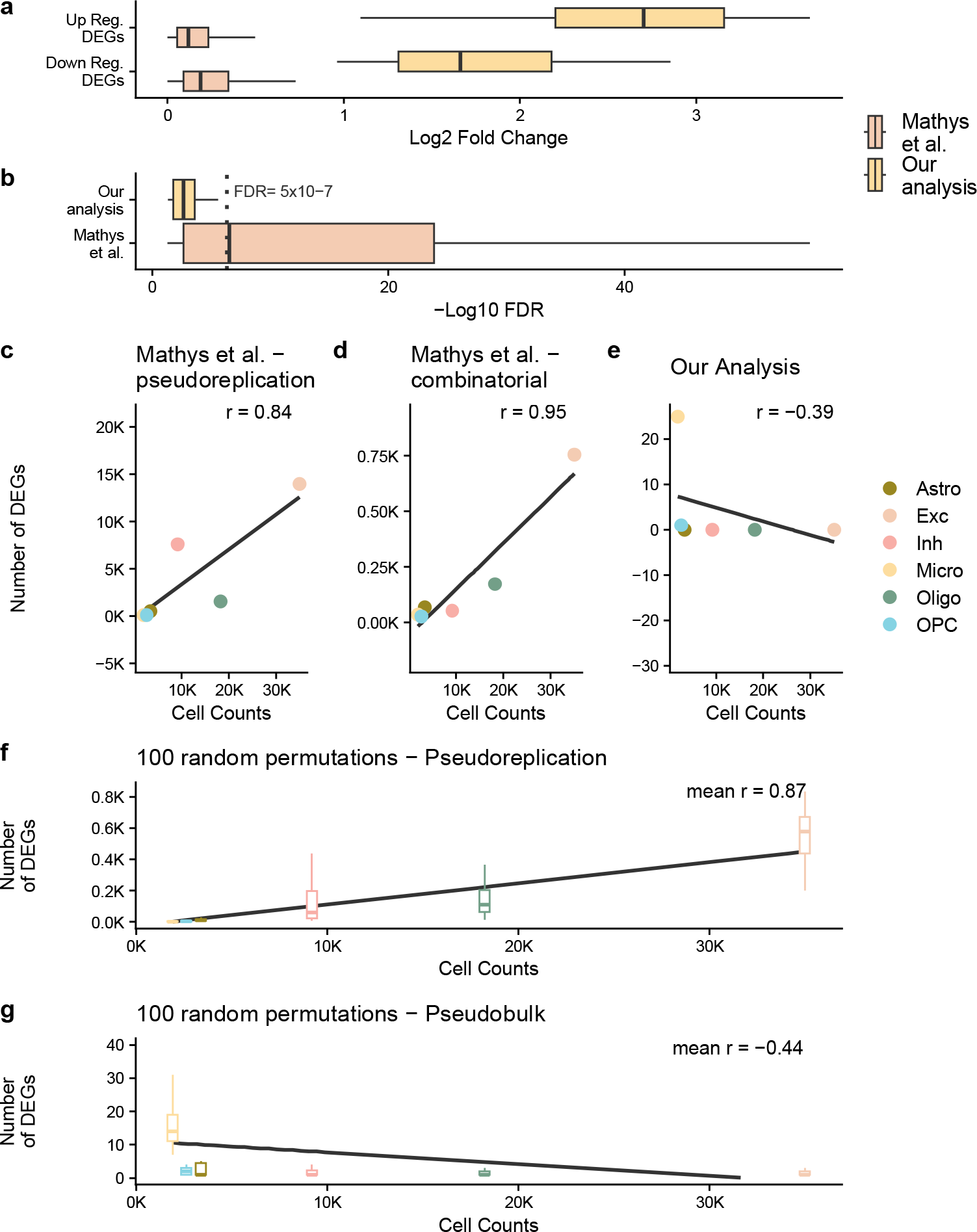
**a, b** highlight the log_2_ fold change and -log_10_ false discovery rate (FDR) of the differentially expressed genes from the author’s original work (Mathys et al.) and our reanalysis (Our analysis). In **b**, we have marked an FDR of 5x10^-7^, dashed grey line, to highlight how small the p-values from Mathys et al.’s analysis are. **c, d, e, f, g** show the Pearson correlation between the cell counts after QC and the number of DEGs identified. For **f, g** analysis, the samples have been randomly mixed between case and control patients. The different cell types are astrocytes (Astro), excitatory neurons (Exc), inhibitory neurons (Inh), microglia (Micro), oligodendrocytes (Oligo) and oligodendrocyte precursor cells (OPC).

A further point which questions the authors’ DE approach is that they identified the vast majority of DEGs in the more abundant, neuronal cell types^1^. However, an increase in the number of cells is not the same as an increase in sample size since these cells are not independent from one another - they come from the same sample. Therefore, an increase in the number of cells should not necessarily result in an increase in the number of DEGs whereas an increase in the number of samples would. This point is the major issue with pseudoreplication approaches which over-estimate confidence when performing differential expression due to the statistical dependence between cells from the same patient not being considered^12,14^. In our opinion, it makes more sense to identify the majority of large effect size DEGs in microglia which recent work has established is the primary cell type by which the genetic risk for Alzheimer’s disease acts^18,19^. This is what we found with our pseudobulk DE approach - 96% of all DEGs were in microglia (Supplementary Table 2) whereas only 3% of the authors’ DEGs were in microglia.

Although it has been proven that pseudoreplication approaches result in false positives by artificially inflating the confidence from non-independent samples, we wanted to investigate the effect of the approach on the authors’ dataset. We ran the same cell-level analysis approach – a Wilcoxon rank-sum test and FDR multiple-testing correction, 100 times whilst randomly permuting the patient identifiers (Fig. 1f). We would expect to find minimal DEGs with this approach given the random mixing of case and control patients. However, this pseudoreplication approach consistently found high numbers of DEGs and we observe the same correlation between the number of cells and number of DEGs as with the authors results. We did not observe the same pattern when running the same analysis with pseudobulk differential expression (Fig. 1g). As a result, we conclude that integrating this pseudoreplication approach with a mixed model like the authors proposed just artificially inflates the test confidence for a random sample of the genes resulting in more false discoveries in cell types with bigger counts.

Up to this point, to compare the effect of the DE approaches in isolation, we analysed the same processed data from the authors as opposed to our reprocessed data. We also performed pseudobulk DE on our reprocessed data and found 16 unique DEGs (Supplementary Table 4). It is worth noting that the fold change correlation between our two DE analyses (reprocessed data vs authors processed data) on the identified DEGs is only moderate (Pearson R of 0.57) and is lower than that of the correlation between pseudoreplication and pseudobulk on the same dataset (Supplementary Table 3). This highlights the effect that the low quality, high mitochondrial read cells have on DE analysis.

## Conclusion

In conclusion, the authors’ analysis has been highly influential in the field with numerous studies undertaken based on their results, something we show has uncertain foundations. However, we would like to highlight that the use of pseudoreplication in neuroscience research is not isolated to the author’s work, others have used this approach^20,21,22^ and their results should be similarly scrutinised. Here, we provide our processed count matrix with metadata and also, the DEGs identified using an independently validated, differential expression approach so that other researchers can use this rich dataset free from spurious nuclei or DEGs. While the number of DEGs found here are significantly lower, much greater confidence can be had that these are AD relevant genes. The low number of DEGs found may also cause concern given the sample size and cost of collection and sequencing of such datasets. However, the increasing number of snRNA-Seq studies being conducted for AD, creates the opportunity to conduct differential meta-analyses to increase power. Further work is required in the field to develop methods to conduct such analysis, integrating studies and accounting for their the heterogeneity, similar to that which has been done for bulk RNA-Seq^23^. Some such approaches have already been made in COVID-19 research which could be leveraged for neurodegenerative disease^24^.

## Data availability

The differentially expressed genes and processed count matrix from the original study are available with their manuscript. The count matrix and metadata from our reprocessing approach are available via the AD Knowledge Portal (https://adknowledgeportal.org). The AD Knowledge Portal is a platform for accessing data, analyses, and tools generated by the Accelerating Medicines Partnership (AMP-AD) Target Discovery Program and other National Institute on Aging (NIA)-supported programs to enable open-science practices and accelerate translational learning. The data, analyses and tools are shared early in the research cycle without a publication embargo on secondary use. Data is available for general research use according to the following requirements for data access and data attribution (https://adknowledgeportal.org/DataAccess/Instructions). For access to content described in this manuscript see: https://doi.org/10.7303/syn51758062.1. All other relevant scripts and data for working with this dataset and supporting the key findings of this study are available within the article and its Supplementary Information files or from our Github repository: https://github.com/neurogenomics/reanalysis_Mathys_2019.

## Code availability

The differential expression analysis pipeline is available at: https://github.com/neurogenomics/reanalysis_Mathys_2019. This is a general use pipeline which can be run for any single-nucleus or single-cell transcriptomic dataset. The config file containing all the parameters used and quality control overview file for the scFlow run is also available in this repository.

## Acknowledgements

This work was supported by a UKDRI Future Leaders Fellowship [grant number MR/T04327X/1] and the UK Dementia Research Institute, which receives its funding from UK DRI Ltd, funded by the UK Medical Research Council, Alzheimer’s Society and Alzheimer’s Research UK. The results published here are in whole or in part based on data obtained from the AD Knowledge Portal (https://adknowledgeportal.org). The data available in the AD Knowledge Portal would not be possible without the participation of research volunteers and the contribution of data by collaborating researchers. Study data were provided by the Rush Alzheimer’s Disease Center, Rush University Medical Center, Chicago. Data collection was supported through funding by NIA grants P30AG10161 (ROS), R01AG15819 (ROSMAP; genomics and RNAseq), R01AG17917 (MAP), R01AG30146, R01AG36836 (RNAseq), U01AG32984 (genomic and whole exome sequencing), U01AG46152, U01AG61356 (ROSMAP AMP-AD, targeted proteomics), U01AG46161 (TMT proteomics), U01AG61356 (whole genome sequencing, targeted proteomics, ROSMAP AMP-AD), the Illinois Department of Public Health (ROSMAP), and the Translational Genomics Research Institute (genomic). Additional phenotypic data can be requested at www.radc.rush.edu.

## Authors’ Contributions

A.E.M and N.G.S jointly conceived and executed the study. N.N.F conducted the quality control and processing of the data and wrote up these steps. A.E.M wrote the manuscript which was reviewed by N.N.F and N.G.S.

