## Supplementary files for "Avoiding false discoveries: Revisiting an Alzheimer’s disease snRNA-Seq dataset"

**Murphy *et al.***

**Supplementary Information and Tables**

The PDF files includes:

Supplementary Note 1

Supplementary Table 1

Supplementary Figure 1

**Supplementary Notes**

Supplementary Note 1. scFlow steps

*1.1 Quality control of snRNAseq data*

The raw single-nucleus RNA-Sequencing data (10.7303/syn18485175) and the ROSMAP metadata (10.7303/syn3157322) were downloaded from <https://www.synapse.org/> upon acquiring appropriate approval. Downstream primary analyses of gene-cell matrices were performed using our scFlow pipeline^1^. To determine ambient RNA profile and distinguish true nuclei from empty droplets, emptyDrops was used with a lower parameter of <100 counts, an alpha cutoff of ≤0.001, and with 10,000 Monte-Carlo iterations^2^. This approach has been recommended as best practice in the literature^3^. Nuclei were then filtered for ≥200 total counts and ≥200 total expressed genes, which was defined as a minimum of 2 counts in at least 3 cells. We excluded any nuclei with total counts or total expressed genes with more than 4 median absolute deviation (MAD) defined by an adaptive thresholding method. Nuclei were excluded if the proportion of counts mapping to mitochondrial genes was more than 10%, as set out in best-practice guidelines^3^. Doublets were identified using the DoubletFinder algorithm, with a doublets-per-thousand-cells increment of 8 cells (recommended by 10X Genomics), a pK value of 0.005^4^. DoubletFinder was shown to be the best overall performing method in a recent benchmark^5^. The aggregated number of cells and proportions dropped at each step is given in Supplementary Table 1. All files from the scflow run, including quality control statistics are available in the Github repository in the scFLow_files folder: <https://github.com/neurogenomics/reanalysis_Mathys_2019>. This includes sample level genes and cells quality control numbers.

*1.2 Integration and clustering*

The linked inference of genomic experimental relationships (LIGER) package was used to calculate integrative factors across samples^6^. LIGER was recently found to be one of the top performing methods for batch-effect correction^7^. LIGER parameters used included: k: 30, lambda: 5.0, thresh: 0.0001, max_iters: 100, knn_k: 20, min_cells: 2, quantiles: 50, nstart: 10, resolution: 1, num_genes: 3000, center: false. Two-dimensional embeddings of the LIGER integrated factors were calculated using the uniform-manifold approximation and projection (UMAP) algorithm with the following parameters: pca_dims: 50, n_neighbours: 35, init: spectral, metric: euclidean, n_epochs: 200, learning_rate: 1, min_dist: 0.4, spread: 0.85, set_op_mix_ratio: 1, local connectivity: 1, repulsion_strength: 1, negative_sample_rate: 5, fast_sgd: false^8^. The Leiden community detection algorithm was used to detect clusters of cells from the 2D UMAP (LIGER) embeddings; a resolution parameter of 0.001 and a k value of 50 was used^9^. This approach has been noted as best practice by a recent review^10^. Automated cell-typing of the detected clusters was performed as previously described using the Expression Weighted Celltype Enrichment (EWCE) algorithm in scFlow against a previously generated cell-type data reference from the Allan Human Brain Atlas^11,12^. The top five marker genes for each automatically annotated cell-type were determined using Monocle 3 and validated against canonical cell-type markers^13^.

**Supplementary Tables**

| **Quality Control Steps** | **Total Cells** | **Percentage** |
| --- | --- | --- |
| Pre-Quality Control | 35,389,440 |  |
| Total failed | 35,337,874 | 99.85% |
| - Minimum Library Size (n<200) | 35,307,281 | 99.77% |
| - Maximum Library Size | 4,742 | 0.01% |
| - Minimum Expressed Genes (n<200) | 35,312,434 | 99.78% |
| - Maximum Library Size/Expressed Genes (MAD>4) | 2,149 | 0.01% |
| - Proportion Mitochondrial Genes (>=0.1) | 1,097,738 | 3.10% |
| - Multiplets (pK = 0.0054) | 581 | 0.00% |
| Total passed | 51,566 | 0.15% |

***Supplementary Table 1****: Overview of the aggregated number of cells across samples removed at each step of the quality control (QC) as part of scFlow. A more detailed explanation of these steps is given in Supplementary Note 1. Note that cells can fail QC for more than one check so only the total failed and Total passed rows will sum to 100%.*

| **Cell** | **logFC** | **logCPM** | **LR** | **PValue** | **adj_pval** | **HGNC** |
| --- | --- | --- | --- | --- | --- | --- |
| Mic | 2.70178913 | 6.99794619 | 26.1418415 | 3.17E-07 | 0.00061349 | ACRBP |
| Mic | 1.48930071 | 8.06240877 | 28.6361217 | 8.73E-08 | 0.00019303 | APOC1 |
| Mic | 1.09327669 | 8.64199769 | 21.5323014 | 3.48E-06 | 0.00336416 | CD81 |
| Mic | -1.4157681 | 7.93884875 | 23.9955467 | 9.66E-07 | 0.00135806 | CD83 |
| Mic | 3.3782727 | 6.86183548 | 32.0804401 | 1.48E-08 | 4.58E-05 | CLEC1B |
| Mic | 2.84072452 | 6.74370542 | 21.7745509 | 3.07E-06 | 0.00316269 | EGF |
| Mic | 2.55769658 | 6.78345087 | 18.0468872 | 2.16E-05 | 0.01699007 | ELOVL7 |
| Mic | -1.2056098 | 8.33197499 | 22.6644045 | 1.93E-06 | 0.00229576 | IFI44L |
| Mic | -1.6616069 | 7.15366639 | 16.4801274 | 4.92E-05 | 0.03306938 | IFI6 |
| Mic | -1.9809425 | 7.00396289 | 17.9180823 | 2.31E-05 | 0.01699007 | IFIT3 |
| Mic | 2.76502672 | 6.72978805 | 20.6543637 | 5.50E-06 | 0.00472825 | ITGA2B |
| Mic | 1.90963403 | 7.01552233 | 16.3200189 | 5.35E-05 | 0.03448474 | MAP1A |
| Mic | -1.8194508 | 8.26208887 | 45.2221008 | 1.76E-11 | 1.36E-07 | NAMPT |
| Mic | 2.0945044 | 7.11048456 | 20.8068524 | 5.08E-06 | 0.00462318 | NEXN |
| Mic | -2.3789762 | 6.93896985 | 22.3912441 | 2.22E-06 | 0.00245752 | NR4A2 |
| Mic | -2.8553462 | 6.73713862 | 22.8029868 | 1.79E-06 | 0.00229576 | NR4A3 |
| Mic | 3.32873829 | 6.84942721 | 30.955327 | 2.64E-08 | 6.81E-05 | PF4 |
| Mic | 3.4213986 | 6.87326383 | 33.2621657 | 8.05E-09 | 3.11E-05 | PKHD1L1 |
| Mic | 3.64525677 | 6.93422174 | 38.661272 | 5.04E-10 | 2.60E-06 | PPBP |
| Mic | 2.30482679 | 8.10570443 | 60.7932697 | 6.34E-15 | 9.81E-11 | PTPRG |
| Mic | -1.0382468 | 8.11450266 | 15.5968273 | 7.84E-05 | 0.04850839 | RORA |
| Mic | 2.54636649 | 6.69202981 | 17.2532606 | 3.27E-05 | 0.02300507 | SDPR |
| Mic | -0.9629617 | 8.8434334 | 17.9319131 | 2.29E-05 | 0.01699007 | SYTL3 |
| Mic | -1.4215374 | 7.99629806 | 25.4736272 | 4.48E-07 | 0.00077092 | TMEM2 |
| Mic | 2.98901596 | 6.77276641 | 24.2100819 | 8.64E-07 | 0.00133637 | TUBB1 |
| Opc | -2.8274718 | 5.03371292 | 22.1334581 | 2.54E-06 | 0.04176231 | EGR1 |

***Supplementary Table 2****: The differentially expressed genes from our reanalysis, using the same processed data the authors used and pseudobulk differential expression approach.*

| **Pseudoreplication adjusted p-value cut-off** | **Number of genes compared** | **Pearson correlation** |
| --- | --- | --- |
| 0.01 | 20152 | 0.8646269 |
| 0.05 | 23903 | 0.8708275 |
| 0.1 | 26382 | 0.8721126 |
| 0.25 | 32117 | 0.8764692 |
| 0.5 | 42022 | 0.8751554 |
| 1 | 84467 | 0.826248 |

***Supplementary Table 3****: Pearson correlation between our pseudobulk differential expression analysis and the author’s pseudoreplication analysis on all genes found to be significant at different adjusted p-value cut-offs from the author’s pseudoreplication analysis.*

| **Cell** | **logFC** | **logCPM** | **LR** | **PValue** | **adj_pval** | **ensembl_id** | **HGNC** |
| --- | --- | --- | --- | --- | --- | --- | --- |
| OPC | -4.1544663 | 4.92100803 | 21.6911445 | 3.20E-06 | 0.04985906 | ENSG00000166573 | GALR1 |
| Astro | -4.5845276 | 4.7965143 | 22.2367847 | 2.41E-06 | 0.037634 | ENSG00000137959 | IFI44L |
| Micro | -3.7616619 | 7.32875316 | 26.8149688 | 2.24E-07 | 0.00077905 | ENSG00000077238 | IL4R |
| Micro | -2.0681446 | 7.88736441 | 17.5929095 | 2.74E-05 | 0.0346187 | ENSG00000105835 | NAMPT |
| Micro | -1.6757556 | 7.58472506 | 19.1736829 | 1.19E-05 | 0.02076348 | ENSG00000118257 | NRP2 |
| Micro | -3.1556403 | 6.85232653 | 19.2064627 | 1.17E-05 | 0.02076348 | ENSG00000135363 | LMO2 |
| Micro | -3.4339265 | 6.9290472 | 19.5975589 | 9.56E-06 | 0.02076348 | ENSG00000138135 | CH25H |
| Micro | -2.8183109 | 6.77500676 | 16.907959 | 3.92E-05 | 0.04550806 | ENSG00000142408 | CACNG8 |
| Micro | 2.90076647 | 8.34560617 | 45.5144266 | 1.52E-11 | 2.11E-07 | ENSG00000144724 | PTPRG |
| Micro | 3.25867589 | 6.91671013 | 16.5519147 | 4.73E-05 | 0.0490155 | ENSG00000163106 | HPGDS |
| Micro | -2.0290905 | 7.12321166 | 16.4746746 | 4.93E-05 | 0.0490155 | ENSG00000171612 | SLC25A33 |
| Micro | -3.4657301 | 6.93307221 | 19.7883301 | 8.65E-06 | 0.02076348 | ENSG00000172243 | CLEC7A |
| Micro | -4.172807 | 7.16813583 | 34.3515807 | 4.60E-09 | 3.20E-05 | ENSG00000174600 | CMKLR1 |
| Micro | -3.1984588 | 6.87310555 | 18.5335889 | 1.67E-05 | 0.0232342 | ENSG00000227531 | RP11-202G18.1 |
| Micro | 3.40562887 | 6.9381703 | 18.5526502 | 1.65E-05 | 0.0232342 | ENSG00000228058 | RP11-552D4.1 |
| Micro | 4.46073301 | 7.66559163 | 29.7716679 | 4.86E-08 | 0.00022549 | ENSG00000253496 | RP11-13N12.1 |

***Supplementary Table 4****: The differentially expressed genes from our reanalysis, using the reprocessed data and pseudobulk differential expression approach.*

**Supplementary Figures**

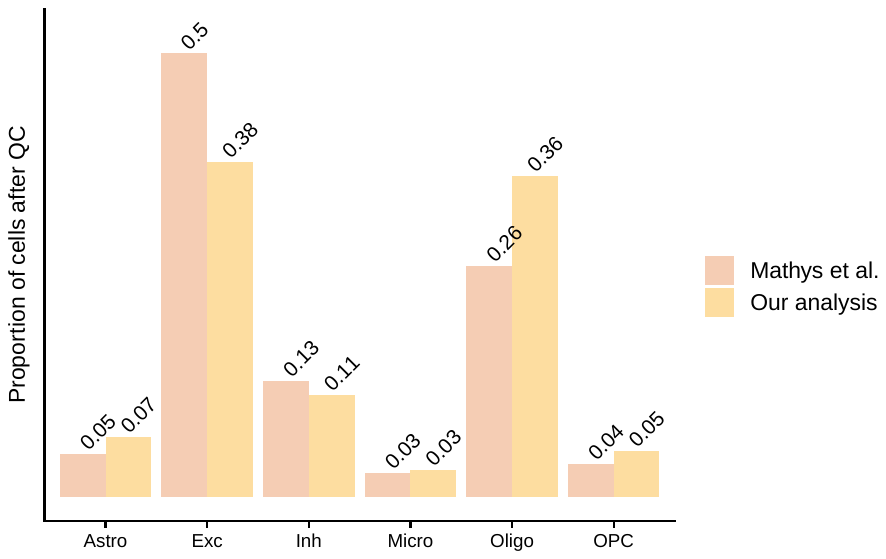

***Supplementary Fig 1****: Shows the proportion of cells left after quality control (QC) from the author’s processing approach (Mathys et al.) and our standardised pipeline approach - scFlow (Our analysis).*

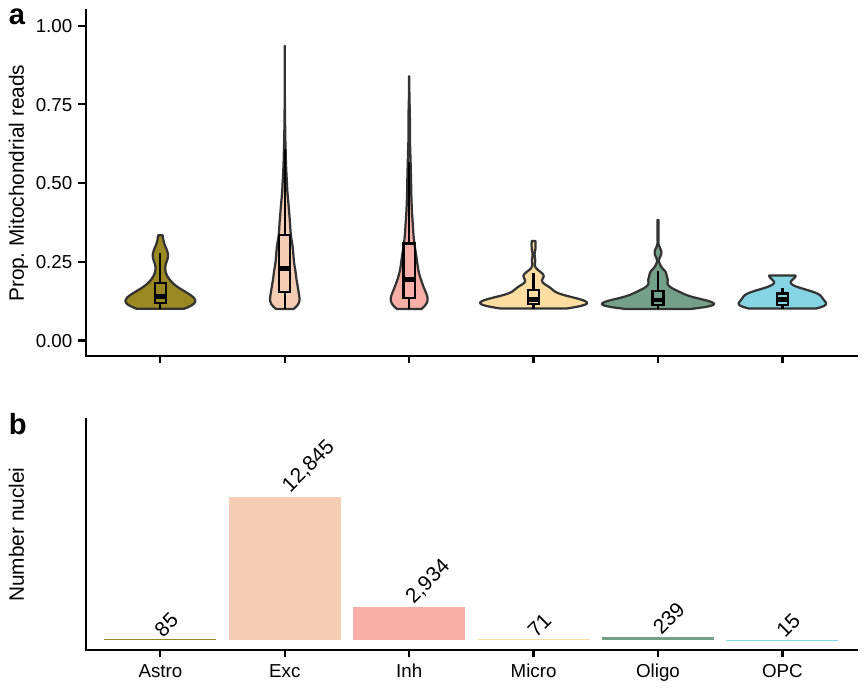

***Supplementary Figure 2****: highlights the nuclei that were removed from our quality control approach, as their proportion of mitochondrial reads were >= 10%, but kept in the authors.* ***a*** *shows the proportion of mitochondrial reads across the different cell types.* ***b*** *gives the number of removed nuclei which were kept by the authors. The different cell types are astrocytes (Ast), excitatory neurons (Ex), inhibitory neurons (In), microglia (Mic), oligodendrocytes (Oli) and oligodendrocyte precursor cells (Opc).*

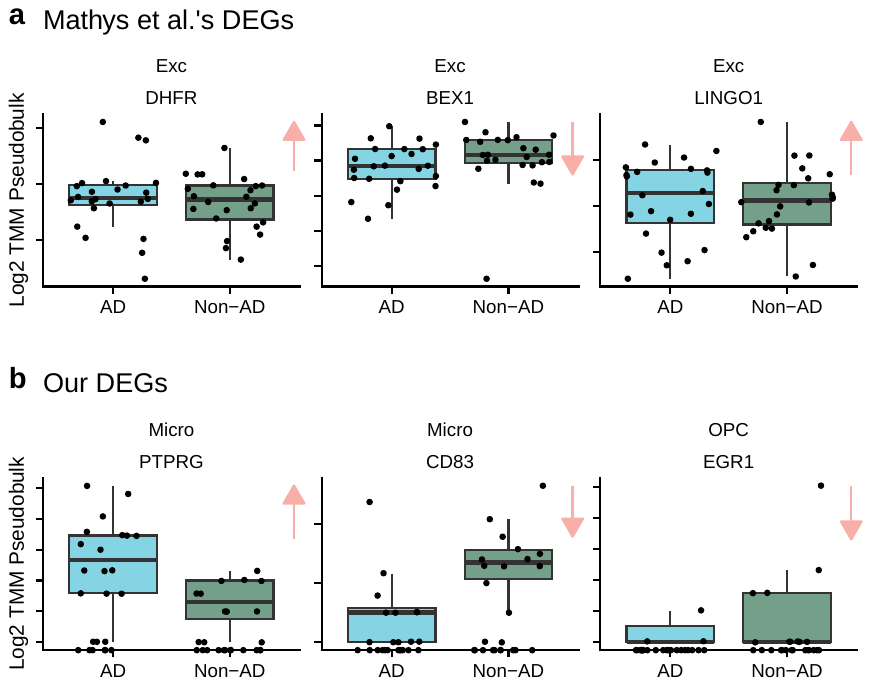

***Supplementary Figure 3****: shows the log_2_ transformed, sum pseudobulked, trimmed mean of M-values (TMM) normalised counts for specific differentially expressed genes. Counts are split by Alzheimer’s disease (AD) pathology and the pink arrow indicates the estimated direction of effect of the DEG in relation to AD.* ***a*** *shows the counts for three neuronal DEGs the authors identified through their differential expression (DE) approach and* ***b*** *shows three DEGs identified by pseudobulk DE analysis (our approach).*
